# Developmental emergence of spatiotemporal coordination in cerebellar Purkinje cell populations

**DOI:** 10.64898/2026.06.12.731562

**Authors:** Kanae Hiyoshi, Kazuhiro Miyanari, Eri Okuda, Narumi Fukuda, Kaito Saito, Tomoya Murayama, Koji Matsuda, Takahisa Matsuzaki, Ryuzo Kawamura, Hiroshi Y. Yoshikawa, Masahiko Hibi, Sachiko Tsuda

## Abstract

Coordinated activity across neuronal populations is fundamental to brain function, yet how such network-level organization emerges during development remains incompletely understood. Here, we performed whole-cerebellar calcium imaging at cellular scale in zebrafish larvae to examine the developmental maturation of Purkinje cell population dynamics. Visual stimulation recruited large, spatially organized populations of Purkinje cells whose responses depended on inferior olive input and were associated with optokinetic behavior. Notably, in the absence of stimuli, Purkinje cells formed transient assemblies exhibiting distance-dependent coordination. During development, long-range coordination progressively emerged, extending initially local correlations into distributed population-wide coordination. Early enucleation, but not dark rearing, disrupted the developmental refinement of long-range coordination and induced aberrant population clustering, suggesting that early retina-dependent signals contribute to cerebellar network development. Together, these findings reveal key organizational features underlying the developmental emergence of coordinated cerebellar population dynamics and suggest that early retina-dependent signals shape population-level organization.

## INTRODUCTION

Coordinated activity among neuronal populations is fundamental to brain function, supporting information processing across neural circuits^1,2^. During development, neural circuits undergo extensive structural and functional remodeling, while patterned population activity is observed in diverse regions, including the retina, spinal cord, cerebral cortex, and cerebellum^2–5^. Such early population activity plays an important role in shaping circuit architecture and functional connectivity^1^. Despite its importance, how coordinated population dynamics emerge and mature across large brain regions during development, particularly at cellular resolution, remains incompletely understood.

The cerebellum provides a powerful system for addressing this question. Purkinje cells, the major output neurons of the cerebellar cortex, integrate excitatory inputs from granule cells and the inferior olive (IO), transforming diverse signals into precisely timed outputs essential for cerebellar computation^6,7^. The cerebellum is characterized by pronounced modular organization at molecular, anatomical, and physiological levels, including striped and patchy compartmentalization patterns^8–11^. Recent studies have shown that Purkinje cell populations encode behavioral variables and exhibit spatially structured activity patterns across the cerebellum, suggesting that population-level coordination is a core component of cerebellar information processing^12–15^. However, the cellular-scale organization of Purkinje cell coordination across the cerebellum and its developmental emergence remain unclear.

To address this gap, zebrafish larvae provide a unique experimental model. Their small size, optical transparency, and genetic accessibility enable large-scale functional imaging of neuronal activity at cellular resolution^16–18^. In addition, the zebrafish cerebellum develops rapidly and shares conserved features of vertebrate cerebellar organization, making it well suited for investigating how Purkinje cell population activity emerges across the cerebellum^19,20^. Previous work in zebrafish has revealed functional regionalization of Purkinje cell populations in developing larvae, with specific cerebellar regions implicated in visual and motor processing^21,22^. Despite these advances, how Purkinje cell population activity becomes coordinated across the developing cerebellum remains unclear. In particular, it remains unknown how Purkinje cell activity is spatially organized at early developmental stages and how this population-level organization changes during cerebellar maturation.

In this study, we conducted whole-cerebellar calcium imaging at cellular scale in zebrafish larvae to examine the developmental emergence of Purkinje cell population dynamics. We identify both stimulus-evoked and resting-state activity patterns, revealing small, transient assemblies of Purkinje cells with distance-dependent coordination as well as larger clusters whose organization progressively matures during development. Manipulation of retinal input further suggests a retina-dependent mechanism that modulates cerebellar network formation. Together, these results reveal the developmental emergence of spatially organized Purkinje cell activity and provide a framework for understanding population-level organization in the cerebellum.

## RESULTS

### Population-level organization of Purkinje cell activity during optokinetic stimulation

To investigate the dynamics of cerebellar Purkinje cell populations during development, we performed wide-field, whole-cerebellar calcium imaging of Purkinje cells using *Tg(aldoca: GCaMP6s)* zebrafish larvae, in which GCaMP6s is selectively expressed in Purkinje cells (Fig.1a, Fig. S1a, Video S1). Immunohistochemical analysis using the Purkinje cell marker Parvalbumin7 confirmed robust expression of GCaMP6s, with approximately 80% of Purkinje cells labeled at 5 days post-fertilization (dpf; Fig. 1b, Fig. S1b; 76.7 ± 4.0% of Parvalbumin7-positive Purkinje cells were GCaMP6s-positive; n = 5 fish, 2352 cells).

**Figure 1.**
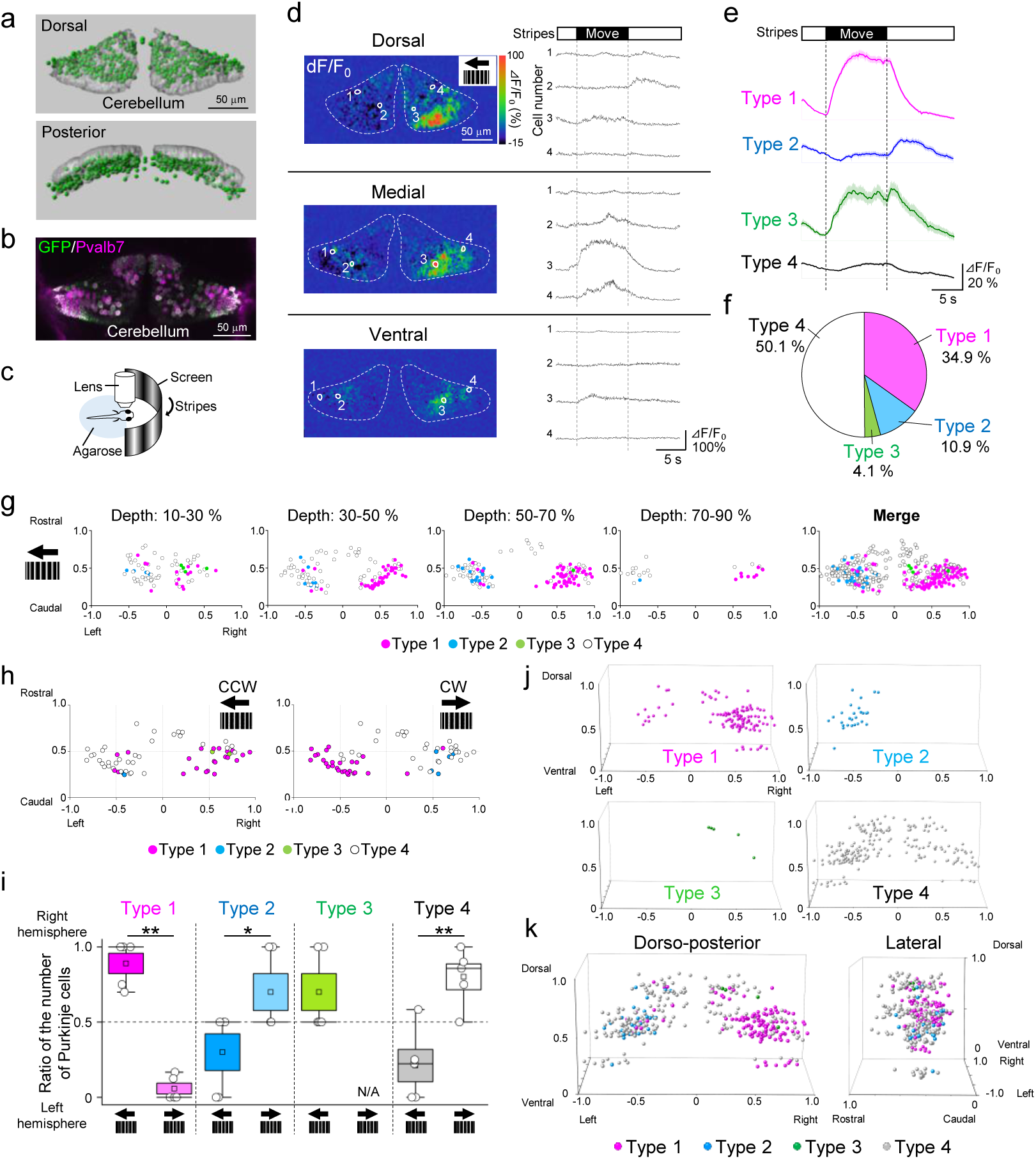
Response patterns and spatial distribution of Purkinje cells during optokinetic response (OKR) stimuli. (a) Spatial distribution of Purkinje cells in the cerebellum of a *Tg(aldoca:GCaMP6s)* zebrafish larva. Spheres indicate the soma positions of Purkinje cells (top: dorsal view, bottom: posterior view). (b) Horizontal cerebellar sections of 5 dpf *Tg(aldoca:GCaMP6s)* larvae stained for GCaMP and Parvalbumin7. (c) Schematic diagram of the OKR system. Larvae were immobilized in agarose and presented with stripe patterns moving either clockwise or counterclockwise. (d) Representative results of Purkinje cell activity at different depths of the cerebellum (dorsal, medial, ventral) during visual stimulation. Changes in fluorescence intensity of GCaMP6s (⊿F/F_0_) during counterclockwise visual stimuli are shown. Regions of interest (ROIs) were located on individual Purkinje cell. (e) Averaged fluorescence changes for four types of Purkinje cell responses (type 1: 394 cells, type 2: 126 cells, type 3: 44 cells, type 4: 580 cells, 10 fish) (f) Ratio of each response type among Purkinje cells (10 fish, type 1: 398 cells, type 2: 124 cells, type 3: 47 cells, type 4: 572 cells) (g) Distribution of Purkinje cell response types (dorsal view, stimulus direction: counterclockwise). Type 1: pink, type 2: blue, type 3: green, type 4: gray. The predominant response type across multiple trials was designated as the representative response type for each Purkinje cell (probability > 0.66, 10 fish, 60 trials, 1141 cells). (h) Distribution of Purkinje cell response types during counterclockwise (CCW) or clockwise (CW) stimuli (5 fish, 62 cells). (i) Laterality of the responsive Purkinje cells changed with stimulus direction. The direction of the stimuli is shown at the bottom (5 fish, *p < 0.05, **p < 0.01, two-sample t-test). (j, k) 3D distribution of the recorded Purkinje cells for each response type in a reference cerebellum (10 fish, 60 trials, 1141 cells).

We next probed Purkinje cell population responses using optokinetic response (OKR), a well-characterized visuomotor behavior that engages cerebellar circuits^23^, and focused on 6 dpf larvae, when both slow and fast phases of OKR were reliably detected (Fig. 1c, Fig. S1c–e). Visual stimuli eliciting OKR evoked Purkinje cell activity across the cerebellum. The spatial pattern of responsive regions depended on stimulus direction (Fig. S1f–h), consistent with previous reports^21,22^.

To resolve response heterogeneity at the cellular level, we next performed high-speed confocal calcium imaging under identical stimulus conditions at multiple cerebellar depths (Fig. 1d, Video S2). Cellular-level analysis revealed multiple response patterns that were classified into four types: type 1 (activated after the onset of stripe movement), type 2 (activated at the offset of stripe movement), type 3 (activated at both onset and offset of stripe movement), and type 4 (exhibited little or no response at either the onset or offset of stripe movement) (Fig. 1e; Fig. S2a). Among Purkinje cells that exhibited evoked responses, type 1 responses were most prevalent, followed by types 2 and 3 (Fig. 1f). These response types exhibited distinct spatial distributions between hemispheres and along dorsoventral axis (Fig. 1g, Fig. S2b–d), with hemispheric asymmetry that depended on stimulus direction (Fig. 1h, i, Fig. S2e, f). Three-dimensional reconstruction revealed that Purkinje cells of each response type formed spatially extended clusters that partially overlapped with other types (Fig. 1j, k).

Because Purkinje cell calcium signals can reflect both sensory and motor-related activity, we next examined the relationship between Purkinje cell activity and fast-phase eye movements, a prominent component of the OKR (Fig. 2a, b). Analysis of calcium transient timing revealed that subsets of Purkinje cells, particularly type 1 and type 4 cells, exhibited activity coincident with the fast phase of OKR (Fig. 2c, Fig. S3a). These fast phase-related Purkinje cells were broadly distributed across the cerebellum, with spatial patterns resembling those observed for visual responses (Fig. 2d, Fig. S3b). To examine the possible function of these responses, we mechanically suppressed eye movements by agarose embedding and reexamined Purkinje cell response profiles (Fig. 2e). With this manipulation, a subset of type 1 cells altered their response profiles (Fig. 2f–h, Fig. S3c, d), suggesting a contribution of sensorimotor integration-related signals. Accordingly, type 1 cells were subdivided into type 1a cells, whose responses were preserved after eye immobilization (sensory-related), and type 1b cells, whose responses were altered by eye immobilization (sensory- and motor-related). Together, these results demonstrate that Purkinje cell activity during OKR is organized into spatially structured populations across the cerebellum (Fig. 2i).

**Figure 2.**
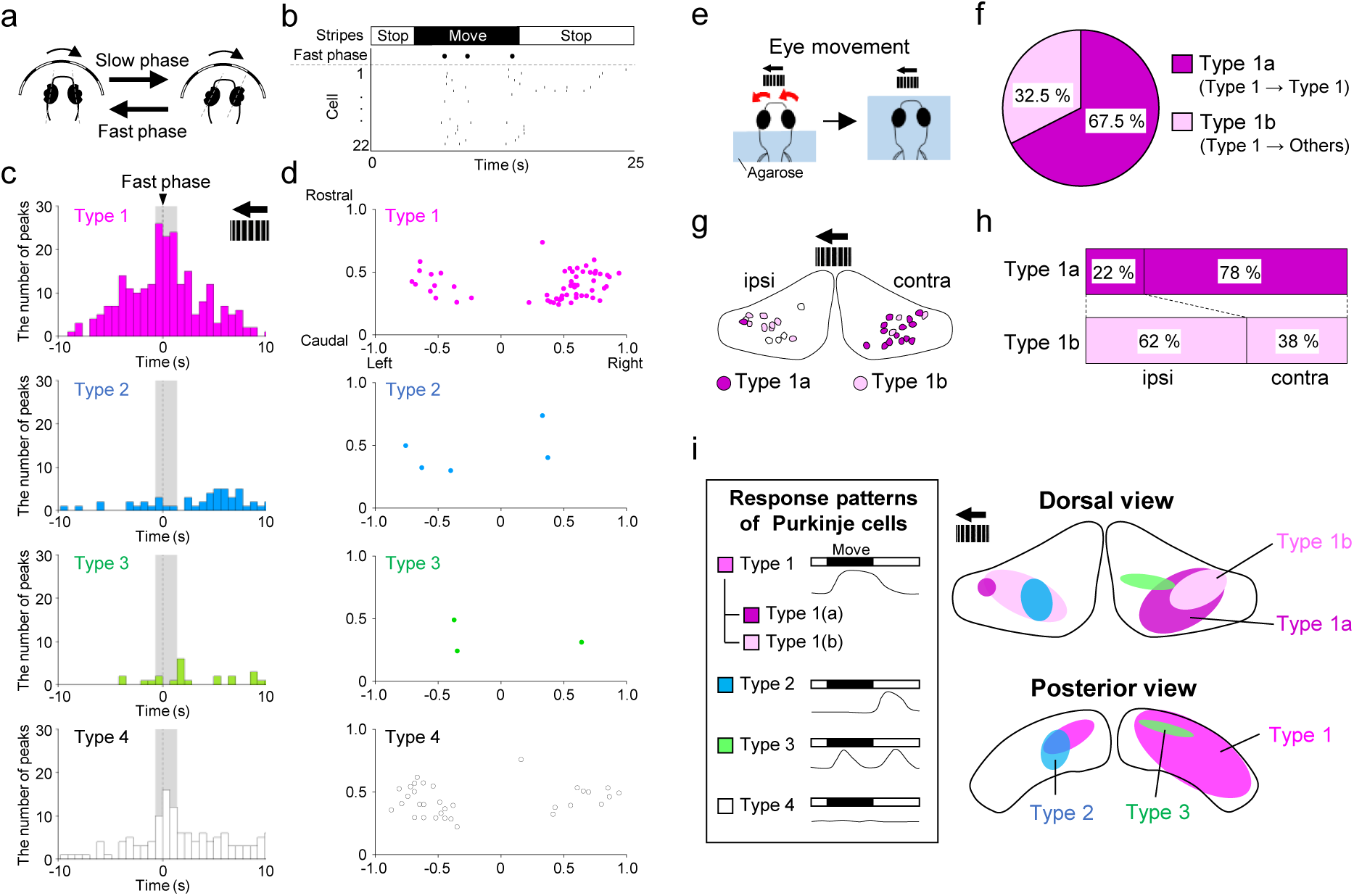
Functional organization of Purkinje cell populations. (a) Schematic diagram of OKR eye movements. (b) Representative time course of fast phases and calcium transients during visual stimulation. (c) Timing of calcium transients in Purkinje cells of each type (7 fish, 12 trials, 484 events). Time zero indicates the timing of the fast phase. (d) Distribution of Purkinje cells with calcium transients coinciding with the fast phase (-0.7 to 1.4 s relative to the fast phase, grey area in (c), 7 fish, 12 trials). (e-h) Blocking eye movements changed the response types of type 1 Purkinje cells. (e) Schematic diagram of the experiments. (f, g) Ratio and distribution of type 1a and 1b Purkinje cells (type 1a: 27 cells, type 1b: 13 cells, 7 fish). (h) Type 1a and 1b Purkinje cells exhibited different distributions between the cerebellar hemispheres (ipsi: ipsilateral hemisphere to the stimulus direction, contra: contralateral hemisphere to the stimulus direction). (i) Distribution of Purkinje cell populations with different response patterns in the cerebellum.

### Inferior olive involvement in Purkinje cell responses to visual stimuli

To understand the mechanisms driving Purkinje cell population activity, we next examined activity in inferior olive (IO) neurons. In contrast to the mammalian cerebellum, calcium transients in zebrafish Purkinje cells are evoked by both IO and granule cell inputs, with the majority of the input originating from the IO^21^. Calcium imaging revealed that subsets of IO neurons were activated immediately following stimulus onset, resembling type 1 Purkinje cell response patterns (Fig. 3a, b, Video S3). Consistent with contralateral olivocerebellar projections^24^, the somata of these IO neurons were predominantly located ipsilateral to the stimulus direction (Fig. 3c). To verify the role of IO neurons in Purkinje cell responses to visual stimuli, we locally ablated IO neurons by ultrashort laser irradiation via two-photon excitation and performed calcium imaging before and after ablation in larvae expressing RFP in IO neurons and GCaMP6s in Purkinje cells (*Tg(hspGFFDMC28C;UAS:RFP;aldoca:GCaMP6s)*) (Fig. 3d). Laser ablation resulted in reduced fluorescence intensity and morphological disruption of IO neurons as we previously demonstrated^25^ (Fig. 3e), and also significantly decreased OKR velocity (Fig. 3f). Moreover, unilateral IO ablation selectively reduced Purkinje cell responses to contralateral visual stimuli (Fig. 3g, h; Fig. S3e). These findings indicate that visually evoked Purkinje cell population activity is shaped by IO input.

**Figure 3.**
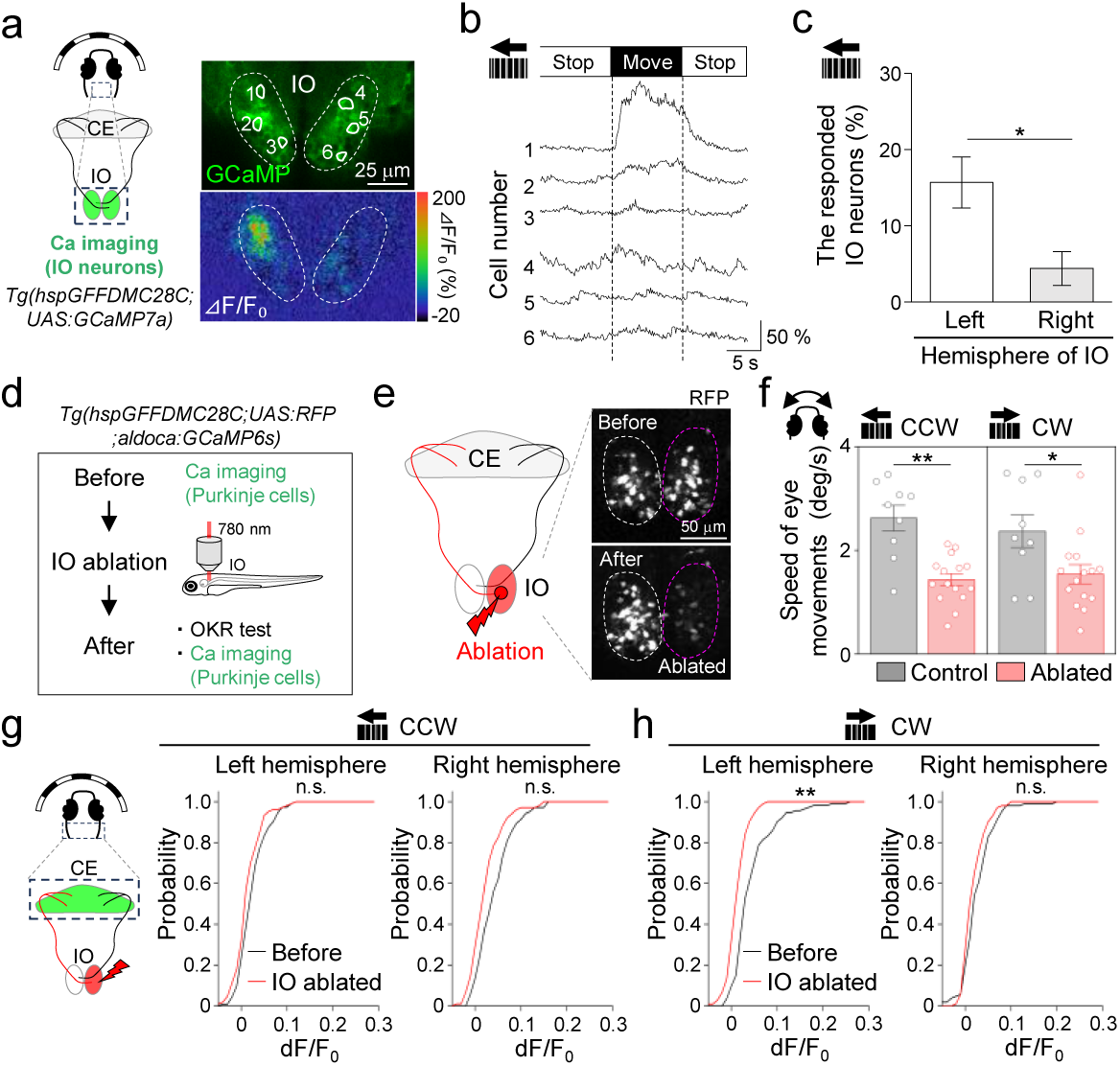
Responses of inferior olive neurons to visual stimulation and their effects on Purkinje cell activity. (a) (Left) Schematic diagram of inferior olive (IO) calcium imaging in response to visual stimuli. (Right) GCaMP7a expression in the IO of *Tg(hspGFFDMC28C;UAS:GCaMP7a)* larvae (top), with a representative image of IO activity during stripe movement (bottom). (b) Fluorescence changes in the IO neurons indicated in (a). (c) IO neurons located in the left hemisphere showed responses to counterclockwise-moving stripes with higher probability (4 fish, *p < 0.05, two-sample t-test). (d) Experimental design of IO ablation. (e) Schematic diagram of IO ablation (left), and stacked images of RFP signals in the IO before and after ablation (right). (f) The speed of slow-phase eye movements in response to visual stimuli was significantly reduced after IO ablation (control: 9 fish, ablated: 15 fish, *p < 0.05, **p < 0.01, two-sample t-test). (g, h) Ablation of the right IO significantly reduced Purkinje cell responses to CW visual stimuli in the cerebellum in the left cerebellar hemisphere. Cumulative probability of maximum activity (⊿F/F_0_) of Purkinje cells during 10 s of stimulation is shown (before: 116 cells, IO ablated: 108 cells, 4 fish, *p < 0.05, **p < 0.01, Kolmogorov–Smirnov test).

### Purkinje cell populations exhibit transient, spatially clustered activity at rest

To determine whether the population-level organization observed during visual stimulation was restricted to stimulus-evoked conditions or reflected a more general feature of cerebellar population dynamics, we next examined Purkinje cell activity under resting conditions in the absence of explicit visual stimulation. Under these conditions, Purkinje cells exhibited activity widely across the cerebellum (Fig. 4a, Video S4), forming populations that varied in position and size over time (Fig. 4a, c, Fig. S4a). We refer to these spatially contiguous active populations as Purkinje cell clusters for further characterization. Cluster area was highly variable, with many clusters being relatively small (mean area: 539.6 ± 690.6 μm²; Fig. 4d, e). Based on area measurements of individual Purkinje cells by single-cell labeling (233.1 ± 13.8 μm²; Fig. 4f, Fig. S4b), we estimated that clusters typically comprised approximately two to three Purkinje cells. Furthermore, as shown in Figure 4g, nearby Purkinje cell clusters often exhibited partial spatial overlaps, suggesting dynamic changes in population activity over time (Fig. 4g, h). These results demonstrate that Purkinje cell population activity in the developing cerebellum is organized into transient and spatially dynamic clusters, rather than fixed assemblies that maintain stable size and position (Fig. 4j). Because mammalian cerebellar modules exhibit a polarized organization along the rostrocaudal axis, as indicated by Zebrin II expression^26–28^, we also examined the orientation of small Purkinje cell clusters during the resting state. Many clusters exhibited a rostrocaudal orientation with a slight lateral inclination (Fig. S4c, d).

**Figure 4.**
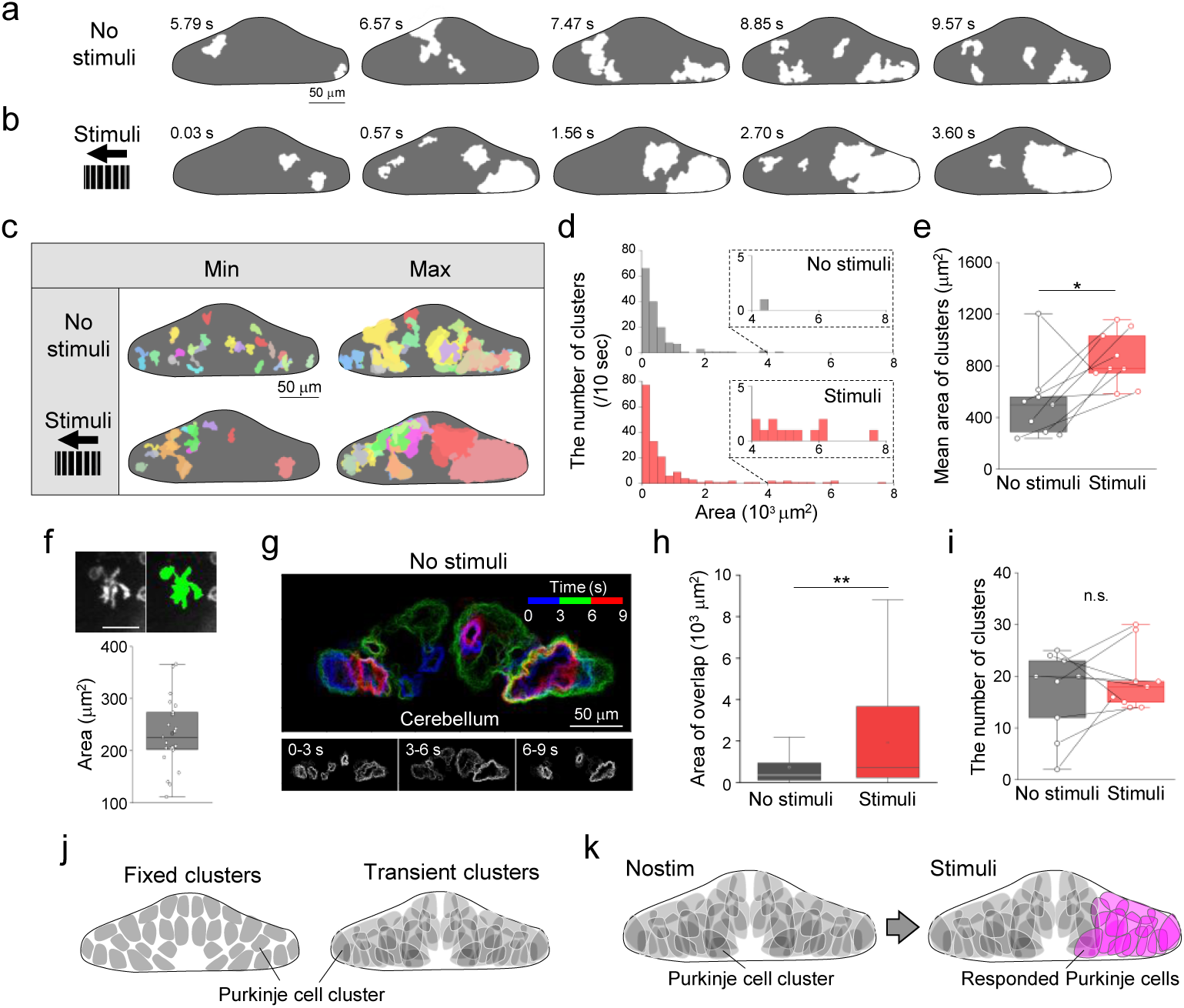
Spatiotemporal dynamics of coordinated Purkinje cell populations. (a, b) Representative images of coordinated activity of Purkinje cells and their changes at resting condition (a) and visual stimulation (b). White regions indicate areas occupied by active Purkinje cell populations. (c) Visualization of the distribution of Purkinje cell clusters at resting condition (no stimuli) and visual stimulation (stimuli). The minimum and maximum area of each Purkinje cell cluster during 10 seconds was overlaid. Each Purkinje cell cluster is shown in a distinct color. The same color indicates the same cluster in both the minimum and maximum panels. (d) Size plots of Purkinje cell clusters under conditions with/without stimuli. The mean area was measured for 10 secs (duration of the visual stimuli). (e) The mean area of Purkinje cell clusters was increased by visual stimuli (no stimuli: 152 clusters, stimuli: 174 clusters, 9 fish, *p < 0.05, two- sample t-test). (f) Representative image of the mosaic-labeled Purkinje cell (top) and the area of labeled Purkinje cells (bottom, 22 cells, 15 fish). (g) Spatiotemporal changes of coordinately activated Purkinje cell populations (Purkinje cell clusters). (Top) The shape of Purkinje cell clusters at three different periods was indicated in blue, green, and red (blue: 0–3 s, green: 3–6 s, red: 6–9 s). Images of each period are shown at the bottom. (h) Overlapped areas between clusters (maximum occupied area for 10 frames) were expanded by visual stimuli (8 fish, **p < 0.01, Mann–Whitney U test). (i) No significant difference was observed in the number of Purkinje cell clusters with/without stimuli (No stimuli: 152 clusters, Stimuli: 174 clusters, 9 fish, two-sample t-test, p = 0.479). (j) Schematic diagrams showing the two possible spatiotemporal dynamics of Purkinje cell clusters. (left) Purkinje cell clusters are made up of fixed populations of Purkinje cells. (right) Purkinje cell clusters are not fixed and are formed with transiently co-activated populations. (k) Expected organization of Purkinje cell clusters in the cerebellum and the changes upon visual stimuli.

Compared with the resting state, visual stimulation significantly increased cluster size (Fig. 4b-e, Fig. S4a) without altering the number of clusters (Fig. 4i). The extent of spatial overlap between Purkinje cell clusters was also significantly increased compared with the resting state (Fig. 4h). Enlarged clusters were preferentially observed in cerebellar regions contralateral to the stimulus, whereas smaller clusters persisted (Fig. 4c). These findings suggest that sensory stimulation recruits coordinated activity across multiple Purkinje cell clusters (Fig. 4k).

### Distance-dependent coordination among Purkinje cells

The transient nature of Purkinje cell clusters described above suggests that Purkinje cells are not independently activated but instead exhibit spatiotemporal coordination. We therefore next asked how such coordination among Purkinje cells is organized across space. To characterize the spatial organization of Purkinje cell coordination, we analyzed pairwise activity correlations as a function of intercellular distance (Fig. 5a, b; Fig. S5a, b). At the population level, coordination among Purkinje cells exhibited distinct spatial patterns depending on their relative positions within or across cerebellar hemispheres. Within each hemisphere, Purkinje cell coordination decreased with increasing intercellular distance (Fig. 5c), whereas Purkinje cell pairs located across hemispheres exhibited elevated coordination over longer distances (Fig. 5d). These results indicate the presence of both local coupling and long-range population-level coordination among Purkinje cells across the cerebellum (Fig. 5g). Consistent with this organization, neighboring Purkinje cells within the same hemisphere exhibited significantly higher activity correlations than more distant pairs, whereas the opposite tendency was observed for interhemispheric pairs (Fig. S5c, d). Cross-correlation analysis further supported this distance-dependent coordination among Purkinje cells (Fig. 5e, f). Moreover, coordination was not uniform across the cerebellum. Along the dorsoventral axis, ventrally located Purkinje cells exhibited significantly higher coordination than dorsally located populations (Fig. S6a-c), while retaining local coupling between Purkinje cells (Fig. S6d).

**Figure 5.**
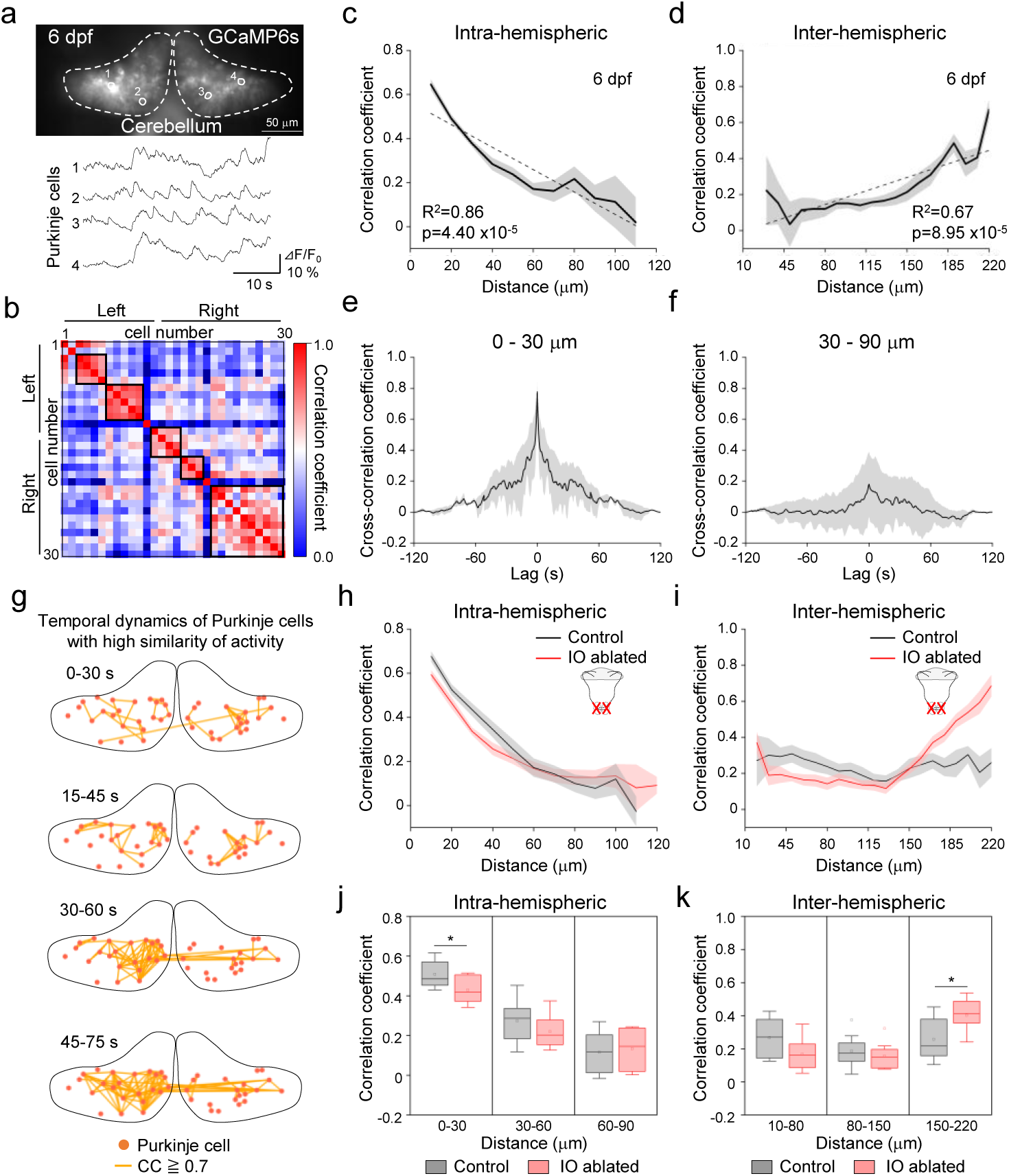
Distance-dependent coordination of Purkinje cell activity in the resting state. (a) Example of Purkinje cell activity in a *Tg(aldoca:GCaMP6s)* larva at 6 dpf. (b) Analysis of Purkinje cell activity similarity based on Pearson correlation. The correlation matrix is aligned using hierarchical clustering (Fig. S5a, b). (c, d). The correlation coefficient of Purkinje cell pairs is plotted against the intercellular distance. Results for Purkinje cell pairs located within or across cerebellar hemispheres are shown in c and d, respectively (n = 7 fish, 3211 intra-hemispheric and 3284 inter-hemispheric pairs). (e, f) Cross-correlation analysis of Purkinje cell activity for pairs with intercellular distances of 0–30 µm (e, 3 fish, 72 pairs) and 30–90 µm (f, 3 fish, 50 pairs). (g) Example of Purkinje cell network structure and its changes (analysis window: 30 s). Purkinje cell pairs with a correlation coefficient greater than 0.7 were visualized with orange lines. Dots indicate the positions of the analyzed Purkinje cells. (h, i) Correlation coefficient of Purkinje cell pairs is plotted against the intercellular distance in control larvae (gray) and larvae with bilateral ablation of the inferior olive (red) (control: 8 fish, 2743 intra-hemispheric pairs and 4211 inter-hemispheric pairs; IO ablated: 10 fish, 2864 intra-hemispheric pairs and 4324 inter-hemispheric pairs). (j, k) Correlation coefficients of Purkinje cell pair activity within (j) and between (k) hemispheres at each intercellular distance (30 µm bins). IO ablation decreased the similarity of Purkinje cell pairs separated by 0–30 µm in the same hemisphere (j) but increased it for pairs separated by 10–80 µm across hemispheres (k) (control: 8 fish, 2742 intra-hemispheric pairs, 4211 inter-hemispheric pairs, IO ablated: 10 fish, 2864 intra-hemispheric pairs, 4324 inter-hemispheric pairs, 10 fish, *p < 0.05, two-sample t-test).

To understand the mechanisms underlying this coordination, we next examined whether olivocerebellar input contributes to the observed distance-dependent coordination patterns. We therefore analyzed Purkinje cell activity following inferior olive (IO) ablation. Following IO ablation, coordinated activity among neighboring Purkinje cells within the same hemisphere (cellular distance: 0–30 μm) was significantly reduced (Fig. 5h, j). In contrast, coordination among distantly located Purkinje cells across hemispheres (cellular distance: 150–220 μm) was significantly increased (Fig. 5i, k). These results indicate that Purkinje cell clusters are associated with distance-dependent coordination patterns that are differentially shaped by olivocerebellar input.

### Developmental emergence of coordinated Purkinje cell activity

To determine how spatial coordination among Purkinje cells emerges during development, we extended our calcium imaging analysis to 3 dpf larvae, shortly after the onset of Purkinje cell differentiation^19,20^, and compared them with 6 dpf larvae. At 3 dpf, fewer Purkinje cells exhibited detectable activity under resting conditions (55.8%, 242 cells, Fig. 6a) compared with 6 dpf larvae, in which nearly all analyzed Purkinje cells were active (97.1%, 309 cells). Within each cerebellar hemisphere, distance-dependent coordination was markedly less pronounced at 3 dpf, although neighboring Purkinje cells still tended to exhibit coordinated activity (Fig. 6b, Fig. S5e). In contrast, coordination among Purkinje cells across hemispheres exhibited a distinct distance-dependent pattern at 3 dpf, with distant pairs showing lower correlations than nearby pairs (Fig. 6c). By 6 dpf, long-range coordination among Purkinje cells across hemispheres increased substantially (Fig. S5f), resulting in the spatial organization characteristic of later developmental stages. These results indicate that local coordination among neighboring Purkinje cells emerges early in development in an immature form, followed by the progressive establishment of long-range inter- hemispheric coordination across the cerebellum.

**Figure 6.**
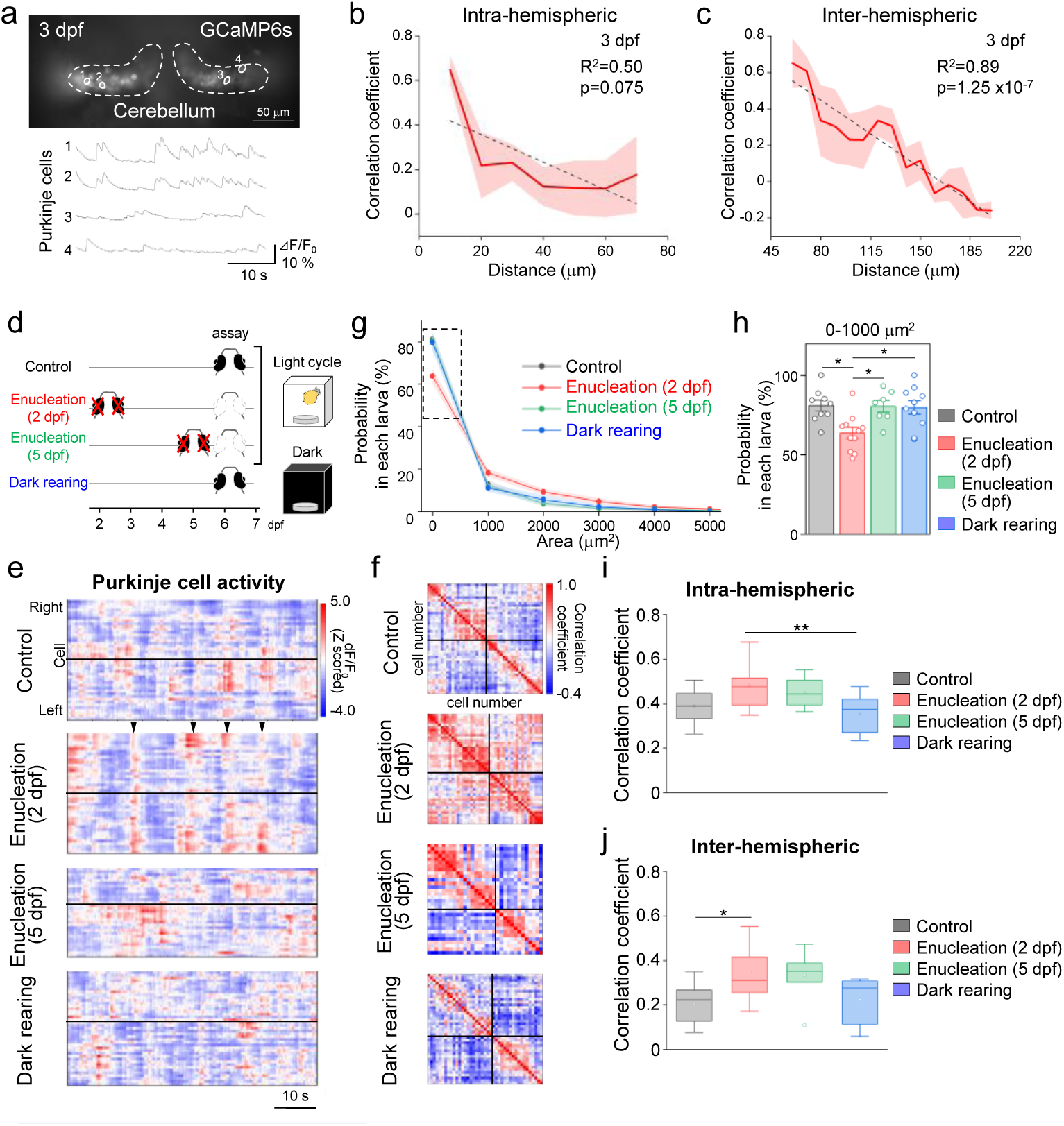
Early retinal input modulates the maturation of Purkinje cell coordination. (a) Example of Purkinje cell activity in a *Tg(aldoca:GCaMP6s)* larva at 3 dpf. (b, c) The correlation coefficient of Purkinje cell pairs is plotted against the intercellular distance. Results for Purkinje cell pairs located within or across cerebellar hemispheres are shown in b and c, respectively (intra-hemispheric pairs: 787, inter-hemispheric pairs: 515, 7 fish). (d) Experimental protocol for the developmental manipulations. (e) Time course of Purkinje cell activity (*Δ*F/F_0_) in the larvae under different conditions in representative larvae (control: 43 cells, enucleation at 2 dpf: 40 cells, enucleation at 5 dpf: 32 cells, dark rearing: 41 cells). (f) Correlation matrix of Purkinje cell activity similarity based on the Pearson correlation in the representative larvae shown in (e). (g) Plots of the area occupied by coordinated Purkinje cell activity (control: 9 fish, enucleation at 2 dpf: 11 fish, enucleation at 5 dpf: 7 fish, Dark rearing: 10 fish). (h) Proportion of coordinated Purkinje cells with small areas (0–1000 µm^2^) was reduced by enucleation at 2 dpf (Control: 9 fish, enucleation at 2 dpf: 11 fish, enucleation at 5 dpf: 7 fish, Dark rearing: 10 fish, *p < 0.05, one-way ANOVA followed by Tukey’s post hoc test). (i, j) Comparison of the correlation coefficient of Purkinje cell activity among four conditions. Results for Purkinje cell pairs located within the same hemispheres and across hemispheres are shown in i and j, respectively (intra-hemispheric pairs, control: 9 fish 2789 pairs, enucleation at 2 dpf: 11 fish, 3792 pairs, enucleation at 5 dpf: 7 fish, 2667 pairs, dark rearing: 4207 pairs, 10 fish; inter-hemispheric pairs, control: 9 fish, 2514 pairs, enucleation at 2 dpf: 11 fish, 3904 pairs, enucleation at 5 dpf: 7 fish, 2741 pairs, dark rearing: 10 fish, 4408 pairs, *p < 0.05, one-way ANOVA followed by Tukey’s post hoc test).

**Figure 7.**
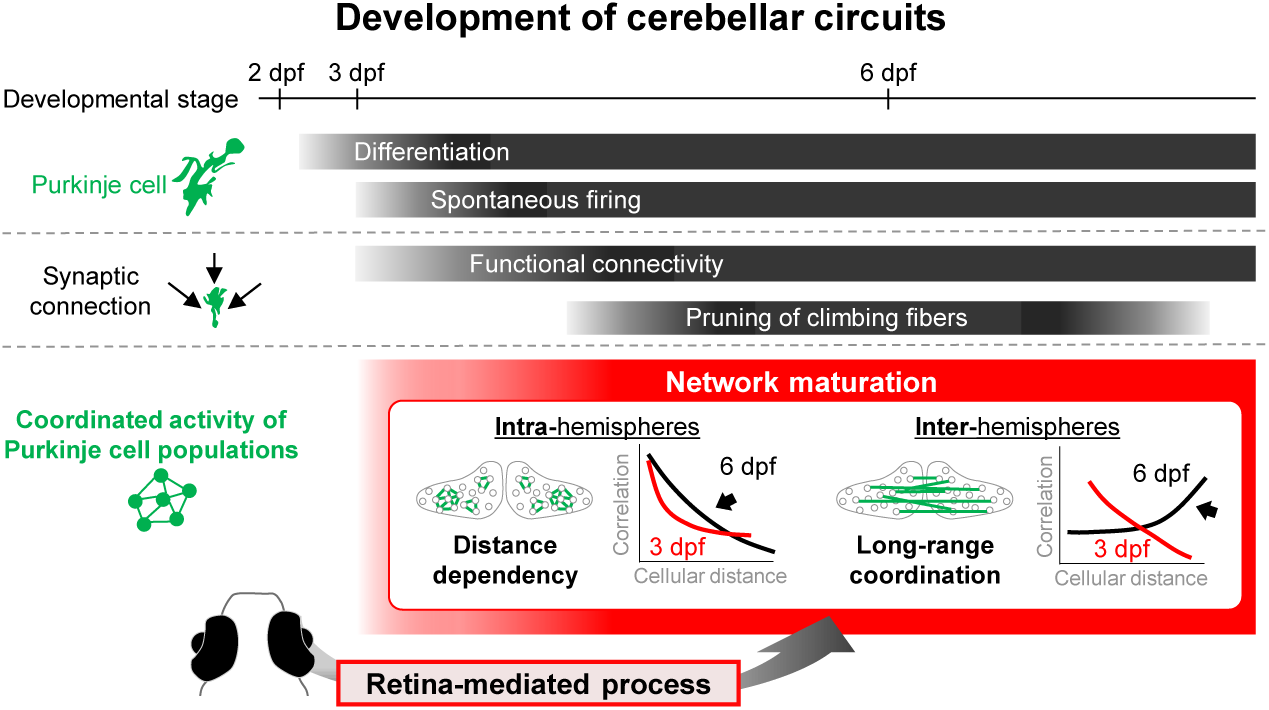
Developmental model of Purkinje cell population coordination. Schematic diagram illustrating cerebellar circuit development from the perspectives of individual Purkinje cell development (differentiation, firing, and synaptic connectivity) and coordinated activity in Purkinje cell populations. The lower panel shows changes in the coordinated activity of Purkinje cells during development.

### Early retina-dependent processes shape the maturation of Purkinje cell population coordination

The developmental reorganization of Purkinje cell coordination between 3 and 6 dpf raised the question of which early signals contribute to this maturation. Because visual stimulation recruited widespread Purkinje cell populations, and visual inputs and retina is known to influence neural circuit refinement during development^1,29,30^, we examined whether the retina contributes to the maturation of Purkinje cell population coordination. To determine whether retinal input before the onset of Purkinje cell differentiation influences the subsequent development of Purkinje cell population activity, we established four experimental groups: enucleation at 2 dpf (before Purkinje cell differentiation), enucleation at 5 dpf (after the initial period of cerebellar circuit assembly), dark-reared larvae, and normally reared controls (Fig. 6d, Fig. S7a). Purkinje cell activity was recorded at 6 dpf under resting conditions.

Early enucleation at 2 dpf markedly altered the spatial organization of coordinated Purkinje cell activity (Fig. 6e). Coordinated activity extended across broader cerebellar regions (Fig. 6e, f) and involved larger Purkinje cell populations (Fig. 6g, h, Fig. S7b, c), prominently across cerebellar hemispheres (Fig. 6i, j). In particular, inter-hemispheric Purkinje cell pairs exhibited elevated coordination at short-to-medium intercellular distances (Fig. S7d, e), resembling the coordination pattern observed at the earlier developmental stage (Fig. 6c). In contrast, enucleation at 5 dpf or dark rearing did not significantly affect Purkinje cell coordination patterns (Fig. 6e–j, Fig. S7b-e). These results suggest that early retina-dependent processes contribute to the maturation and spatial refinement of long-range coordination among Purkinje cell populations across the cerebellum.

## DISCUSSION

### Optical analysis of cerebellar population dynamics at cellular and whole-cerebellar scales

In this study, we investigated the spatiotemporal organization of Purkinje cell population activity in the developing cerebellum, addressing the challenge of linking cellular-level dynamics to large-scale network organization. Although the cerebellum has been viewed as a highly modular structure based on its repetitive anatomical organization^9^, recent imaging studies have revealed distributed activity patterns spanning broad cerebellar regions in mammals and teleosts^14,21,22^. However, how coordinated activity emerges and is organized across the developing cerebellum has remained incompletely understood.

Whole-cerebellar calcium imaging in larval zebrafish enabled us to analyze Purkinje cell activity across the cerebellum while retaining cellular-level spatial information. During optokinetic behavior, we identified multiple evoked response types in Purkinje cells, revealing populations that were spatially distributed across the cerebellum rather than confined to sharply delineated regions. The predominant response type was largely driven by inferior olive input and showed response patterns consistent with previous studies^21,22^, whereas other response types emerged at stimulus offset and may contribute to the temporal regulation of visuomotor responses. Together, these analyses reveal heterogeneous and partially intermingled functional organization in the zebrafish cerebellum, providing a basis for understanding Purkinje cell population dynamics during development.

### Spatial organization and transient coordination of Purkinje cell populations

Our analyses revealed that Purkinje cell population activity is not randomly distributed across the developing cerebellum but instead exhibits a distinct spatial organization. Under resting conditions, Purkinje cells formed transient, spatially clustered populations that continuously changed in size and location rather than remaining as fixed assemblies. Based on area measurements, individual clusters were estimated to comprise two to three Purkinje cells. These clusters were considerably smaller than the Purkinje cell clusters recently reported in the rodent cerebellum, which contain approximately 100 cells^14^. This difference may partly reflect the smaller number of Purkinje cells in the teleost cerebellum^19^, as well as differences in developmental stage.

This spatial organization was further characterized by distance-dependent coordination across multiple spatial scales. Activity correlations declined with increasing distance within cerebellar hemispheres, whereas distant Purkinje cell pairs located across hemispheres exhibited elevated coordination. Thus, cerebellar population coordination exhibited a spatial organization characterized by local coupling together with long-range inter-hemispheric coordination. Similar distance-dependent coordination has been reported within local Purkinje cell populations in the mature mammalian cerebellum^14,31,32^, where shared inferior olive input is considered an important contributor. Consistent with this view, IO ablation reduced coordination among neighboring intra-hemispheric Purkinje cells while increasing coordination among distant inter-hemispheric pairs. These findings suggest that olivocerebellar input shapes the spatial organization of Purkinje cell coordination rather than simply enhancing synchronization globally. Together, our findings extend this principle of distance-dependent coordination from local Purkinje cell populations to the entire developing vertebrate cerebellum and provide a spatial framework for understanding Purkinje cell population dynamics during development.

Visual stimulation further expanded the size and spatial overlap of Purkinje cell activity clusters, consistent with recent studies showing that sensory stimulation recruits distributed Purkinje cell populations across broader cerebellar regions than predicted from localized sensory representations^14,21^. Coordination strength also differed along the dorsoventral axis, with ventral Purkinje cells exhibiting stronger coordination than dorsal populations. Because dorsal and ventral Purkinje cells project to distinct downstream targets in zebrafish^21,22,33^, these differences may reflect functional specializations within cerebellar input–output pathways^34^. Future studies simultaneously examining Purkinje cells and downstream targets, including eurydendroid cells, the teleost homologs of mammalian deep cerebellar nuclei neurons, will clarify how this spatial organization is transformed into cerebellar output.

The spatial organization identified in this study may have implications for understanding cerebellar computation. Purkinje cell synchrony is known to regulate the timing of cerebellar nuclear output^35^, highlighting the functional relevance of population-level temporal organization. Dynamic patterns of Purkinje cell population activity may expand the repertoire of cerebellar population states and thereby support flexible cerebellar output. Integrating large-scale activity measurements with circuit manipulations and computational models will be important for examining this possibility.

### Developmental emergence of local and long-range cerebellar coordination

A major finding of this study is that coordinated Purkinje cell activity emerges progressively across development. Shortly after the onset of Purkinje cell differentiation, distance-dependent coordination within each hemisphere was still weak, although neighboring Purkinje cells tended to exhibit coordinated activity. This finding suggests that local coupling is established early but initially remains immature. In contrast, long-range coordination, particularly between Purkinje cells located across cerebellar hemispheres, became prominent at later developmental stages. These observations suggest that cerebellar population organization develops through the progressive expansion of coordination from local to long-range spatial scales.

In mammals, Purkinje cells exhibit high local synchrony during early postnatal development, which decreases as climbing fiber innervation is refined ^5^. In contrast, our whole-cerebellar analyses in zebrafish revealed the emergence of coordination across broader spatial scales. This apparently different developmental profiles may partly reflect the distinct spatial scales captured by the two experimental approaches. In addition, mammalian Purkinje cell calcium imaging predominantly reflects climbing-fiber-driven complex spikes, whereas Purkinje cell calcium signals in zebrafish contain both complex-spike- and simple-spike-related components^21^. Accordingly, our analyses may capture broader aspects of cerebellar population coordination beyond climbing-fiber synchrony alone.

In zebrafish, Purkinje cell differentiation begins around 3 dpf^19,33^, whereas functional climbing fiber input matures later^36^. Considering this rapid developmental timeline, the coordination patterns observed at early developmental stages likely reflect immature phases of cerebellar circuit assembly. Notably, long-range coordinated activity became prominent between Purkinje cells located across cerebellar hemispheres at 6 dpf, suggesting rapid reorganization of cerebellar functional organization during larval development. The mechanisms underlying early local coordination remain unclear. Because functional input from climbing fibers and granule cells is still limited^36^, electrical coupling^37^ or correlated intrinsic activity may also contribute to the early coordination among neighboring Purkinje cells. These observations support a developmental model in which early local coordination provides a scaffold for the subsequent emergence of long- range, spatially organized cerebellar population coordination. As development proceeds, the integration of extracerebellar inputs and the refinement of synaptic connectivity are likely to promote this transition.

### Retina-dependent modulation of cerebellar network maturation

Our perturbation experiments suggest that early retina-derived signals modulate the developmental maturation of cerebellar coordination. Enucleation at 2 dpf, before Purkinje cell differentiation and early cerebellar circuit assembly, altered Purkinje cell coordination at 6 dpf, producing a distance-dependent pattern resembling that observed at an earlier developmental stage, with elevated coordination at short-to-medium inter-hemispheric distances. In contrast, neither enucleation at 5 dpf nor dark rearing produced comparable effects. Because visual processing has not matured by 2 dpf in zebrafish^38^, these findings suggest that retina-dependent developmental processes, rather than visual experience itself, contribute to cerebellar network assembly. This raises the possibility that spontaneous retinal activity or other early retina-derived signals influence cerebellar maturation, a possibility that will require direct testing in future studies.

More broadly, these findings suggest that early retinal influences contribute to the developmental refinement of cerebellar population coordination. Notably, enucleated larvae exhibited coordination patterns resembling those observed in younger control larvae, raising the possibility that early retinal perturbation interferes with the transition from immature coordination patterns to spatially organized long-range coordination. Further studies examining retinal activity together with cerebellar circuit development will determine how early retina signals contribute to the formation of cerebellar networks during development. Because our analyses were based on calcium transients, the precise temporal synchrony underlying Purkinje cell coordination remains unresolved. The application of genetically encoded voltage indicators in zebrafish^39–41^ may provide further insight into the temporal precision and network mechanisms underlying the spatiotemporal dynamics of Purkinje cell populations.

Together, our findings identify fundamental organizational features underlying the developmental emergence of coordinated cerebellar population dynamics. By combining whole-cerebellar imaging with developmental and perturbation analyses, this study provides a framework for investigating how coordinated neuronal populations emerge during brain development.

## ACKNOWLEDGMENTS

We thank Koichi Kawakami for providing fish lines and constructs, and Junichi Nakai, Masamichi Ohkura, Keiko Ando, and Kyo Yamasu for their technical support and helpful discussions. This work was partly supported by JSPS KAKENHI Grant Numbers JP15K20907 and JP19K06756 to S.T., a Grant-in-Aid for JSPS Fellows (JP21J11883) to K.H., and the JST FOREST Program (JPMJFR205N) to T.M.

## AUTHOR CONTRIBUTIONS

K.H. and S.T. conceived and designed the experiments. K.H., K.M., E.O., N.F., K.S., T.M., T.M., R.K., H.Y., M.H., and S.T. performed the experiments and analyses. K.H., K.M., and S.T. wrote the manuscript. All authors reviewed the manuscript.

## DECLARATION OF INTERESTS

The authors declare no competing interests.

## SUPPLEMENTAL INFORMATION

**Figure S1. Development of OKR and response of Purkinje cell populations to visual stimuli**

(a) GCaMP6s signals in the cerebellum of a *Tg(aldoca:GCaMP6s)* larva. (b) Horizontal cerebellar sections of *Tg(aldoca:GCaMP6s)* larvae labeled by GCaMP (top left) and Parvalbumin7 (top right) at 5 dpf. Magnified images were shown at the bottom. Merged low magnification images of both signals are shown in Fig. 1b. (c) Representative eye movement traces during the visual stimuli at 3, 6, and 8 dpf. The dots indicate the timing of the fast phase. (d, e) Speed of the slow phase (d) and the number of fast phases (e) during stripe movement (10 s). Tukey-kramer, *p < 0.05, 5 fish (3 dpf), 5 fish (6 dpf), 4 fish (8 dpf). (f) (Top) Representative image showing fluorescence changes of GaMP6s during the visual stimuli. (Bottom) Traces of the eye movements and fluorescence changes in the left and right hemispheres during the visual stimuli. (g, h) Peak amplitude of fluorescence changes in each hemisphere during stripes moving (stimulus direction: CW (g), CCW (h), two-sample t-test, *p < 0.05, 3 fish).

**Figure S2. Distribution of Purkinje cells in the cerebellum in response to visual stimuli**

(a) The time distribution of calcium transients for each Purkinje response type (7 fish, 9 planes, 12 trials, 484 transients). (b) Standardization method of Purkinje cell location in the cerebellum (left: XY axis, right: Z axis). (c) Depth of recorded planes along the dorso-ventral axis (11.4 – 88.9%, 12 fish). (d) Number of Purkinje cells in each response type at four different cerebellar depths (10 fish, 1141 cells, Steel-Dwass, *: p < 0.05, **: p < 0.01). (e, f) The number of Purkinje cells of each type along the left-right (e) and dorso-ventral (f) axis (10 fish, *p < 0.05, **p < 0.01, (e) two-sample t-test, (f) Steel-Dwass). Error bars indicate standard error.

**Figure S3. Purkinje cell activity changes associated with eye movement suppression and inferior olive ablation**

(a) Timing of calcium transients of Purkinje cells in relation to eye movements. Time zero indicates the fast phase (7 fish, 172 cells, 484 events). All four response types were combined in this histogram. (b) Distribution of Purkinje cells exhibiting calcium transients during the fast phase, shown in gray in (a). (c, d) Examples of Purkinje cell activity in response to visual stimuli with (c) and without (d) eye movement. (e) Responses of representative Purkinje cells to visual stimuli before (left) and after (right) inferior olive ablation. The right hemisphere of the inferior olive was ablated in the *Tg(hspGFFDMC28C; UAS: RFP; aldoca: GCaMP6s)* larva.

**Figure S4. Spatiotemporal changes in activated Purkinje cell populations**

(a) Minimum and maximum size of Purkinje cell clusters in 10 secs. (b) Representative images of Purkinje cells labeled by *pT2K-aldoca-gap43-venus* pDNA injection. The left panel shows the respective Purkinje cell binarized by fluorescence intensity. The right panels represent confocal microscopy images. (c, d) The orientation of Purkinje cell clusters observed during the 10-second resting state. The orientation of ellipse-fitted objects was evaluated as a morphological feature based on their major axis direction. (c) Comparison of the number of horizontally and vertically oriented objects (8 fish, paired two-sample t-test, p=0.15036). (d) Orientation distribution of objects on each side of the cerebellar hemisphere (gray: right hemisphere, white: left hemisphere, 4 fish).

**Figure S5. Similarities of Purkinje cell activities in the resting state in 3 and 6 dpf larvae**

(a) Representative dendrogram showing the similarity of Purkinje cell activities based on hierarchical clustering. (b) Representative distribution of Purkinje cell populations exhibiting high correlations of activity (Pearson distance less than 0.25, shown in (a)) is visualized in the cerebellum. (c, d) Similarity of activity between Purkinje cell pairs depending on the intercellular distance in the intra- (c) and inter-hemispheres (d) in 6 dpf larvae (6 dpf: 7 fish, *p < 0.05, **p < 0.01, one-way ANOVA followed by Tukey’s post hoc test). (e, f) Comparisons of the similarity in activity between Purkinje cell pairs in 3 dpf and 6 dpf larvae (3 dpf: 7 fish, 6 dpf: 7 fish, **p < 0.01, two-sample t-test).

**Figure S6. Dorsoventral organization of the cerebellum and similarity of Purkinje cell activity**

(a) Distribution of Purkinje cells across cerebellar depths (top) and correlation matrices of Purkinje cell pairs at each depth (bottom). Depth levels: 10.5% (dorsal), 26.3% (medial), and 50.0% (ventral). (b-d) Similarity of activity among intra-hemispheric Purkinje cell pairs at different cerebellar depths (4 fish, one-way ANOVA followed by Tukey’s post hoc test, * p<0.05). (b) Dorsoventral comparison across all analyzed Purkinje cell pairs. (c, d) Relationship between intercellular distance and activity similarity.

**Figure S7. Changes in Purkinje cell activity following retinal input perturbations during cerebellar development**

(a) Representative images of enucleated larva. (b) Cumulative probability of the size of collectively activated Purkinje cell populations across all analyzed larvae. (c) Size distribution of collectively activated Purkinje cell populations, binned in 1000 μm^2^ (Control: 9 fish, enucleation at 2dpf: 11 fish, enucleation at 5dpf: 7 fish, Dark rearing: 10 fish). Merged plot is shown in Fig. 6f. (d, e) Correlation coefficients of Purkinje cell activity for all intra-hemispheric (d) and inter-hemispheric (e) pairs (Control: 9 fish; enucleation at 2dpf: 11 fish, enucleation at 5dpf: 7 fish; dark rearing: 10 fish, one-way ANOVA followed by Tukey’s post hoc test).

**Video S1. Purkinje cell distribution in the cerebellum of a *Tg(aldoca:GCaMP6s)* larva (related to Figure 1a, b)**

The distribution of GCaMP-expressing Purkinje cell somata is shown as spheres.

**Video S2. Purkinje cell responses to visual stimuli at different depths in the cerebellum (related to Figure 1d)**

Purkinje cell activity in the dorsal, medial, and ventral cerebellum of the same *Tg(aldoca: GCaMP6s)* larva. Changes in fluorescence intensity of GCaMP6s (*Δ*F/F_0_) during counterclockwise visual stimuli are shown.

**Video S3. Response of inferior olive neurons to visual stimulation (related to Figure 3a)**

Representative calcium imaging data in the inferior olive of *Tg(hspGFFDMC28C; UAS: GCaMP7a)* larvae during visual stimulation. The size is shown in pseudo-color using ⊿F/F_0_, with the color scale shown in Figure 3a.

**Video S4. Spatiotemporal transitions of activated Purkinje cell populations (related to Figure 4a)**

Background shows representative Purkinje cell activity (*Δ*F/F_0_) under resting conditions. To visualize coordinated Purkinje cell activities and their changes, Purkinje cell populations activated above a threshold are masked in white.

## METHOD DETAILS

### Fish lines

Wild-type zebrafish (*Danio rerio*) with an RW genetic background, *Tg(aldoca: GCaMP6s)* transgenic fish, and *Tg(hspGFFDMC28C; UAS: EGFP)* transgenic fish (Takeuchi et al. 2015) were used for the experiments. To generate the *Tg(aldoca:GCaMP6s)* strain, the entry vectors pENTR-L1-aldoca-R5 ^24^ and pENTR-L5-GCaMP6s-SV40pAS-L2, containing the GCaMP6s cDNA and SV40 polyadenylation signal derived from pGP-CMV-GCaMP6s (Addgene #40753) ^42^, were used for Gateway-mediated recombination into the Tol2 destination vector pT2KDEST-RfaF ^43^. A mixture containing 25 pg of the Tol2 plasmid and 25 pg of transposase-capped RNA was injected into one-cell-stage embryos. Transgene expression was confirmed by GCaMP6s fluorescence. All fish lines were provided by the National Bioresource Project (NBRP). Adult zebrafish were maintained in an incubator with a 14 h light/10 h dark cycle. For most experiments, larvae were incubated at 28 °C under illumination greater than 100 lux. For dark rearing, some larvae were placed in a dark incubator (0 lux) after fertilization. All experiments involving live fish were approved by the Committee for Animal Care and Use of Saitama University.

### Optokinetic response test

Zebrafish larvae were anesthetized in 0.02% tricaine (Ethyl 3-aminobenzoate methanesulfonate salt, Sigma-Aldrich) for 15 min and washed with external solution (134 mM NaCl, 2.9 mM KCl, 2.1 mM CaCl2, 1.2 mM MgCl2, 10 mM glucose, 10 mM HEPES). The larvae were then mounted on 2% low-melting-point agarose (Sigma) in the center of a 35 mm dish, with agarose surrounding the eyes removed. For experiments involving mechanical suppression of eye movements, larvae were carefully mounted to minimize eye movements, and the agarose surrounding the eyes was not removed. To minimize body movements, the tail was positioned in a U-shape, and the dish was filled with 1% methylcellulose (Methyl Cellulose 4000, Wako, 136-02155) or 1/3 Ringer solution (1x Ringer solution: NaCl 116 mM, KCl 2.9 mM, CaCl2 1.8 mM, HEPES 5 mM). To induce OKR, the horizontally moving stripes (duration: 10 s, velocity: 30.3 degrees/s) were presented. These movies were projected onto a white-column screen using a mini-projector (ASUS E1/Vivitek QUMI Q1). Video recordings of the entire experimental system were also made to obtain the timing information of the visual stimuli (120 fps, iPhone 6, Apple). Eye movements were imaged with a wide-field microscope (FN-1, Plan Apo λ 10×/0.45 NA lens, NIKON) equipped with a cMOS camera (ORCA-Flash 4.0, Hamamatsu photonics, HC Image, 33.3-49.9 fps).

### Quantitative analysis of eye movement

Eye positions were tracked by detecting the eye region using image thresholding and elliptical approximations. To calculate the frequency of the fast phase in OKR, after eye movements along the stripe movement direction, a clear reversal movement (a change of 2 °or more in the reversed direction) was regarded as the fast phase, and the number of events was counted. The velocity of the slow phase was analyzed as the angular velocity of the eye during the stimulus (stripe movement), excluding fast phases.

To combine eye movement recording with calcium imaging, eye movements were recorded using an infrared cMOS camera (Blackfly S, BFS-U3-32S4M-C, Point Grey) located beneath the microscope stage and controlled with the SpinView program (Point Grey, 60 or 133 fps). For this specimen, an LED light (850 nm, LDL-74X27IR2-850, CCS) was used for illumination. The camera was equipped with an infrared (IR) transmission filter (IR 76, FUJIFILM). The acquisition timing of the calcium and behavior recordings were detected using a DAQ interface (USB-6212 BNC; National Instruments) with WinWCP software (University of Strathclyde, Scotland, UK).

### Immunostaining and cell counting

Whole-mount immunostaining of zebrafish larvae (*Tg(aldoca:GCaMP6s)*, 5 dpf) was performed as described previously ^25^. In brief, anti-Parvalbumin7 antibody (mouse, 1:1000) and anti-GFP antibody (rat, 1:1000) were used as primary antibodies, and Alexa Fluor 555 goat anti-mouse IgG antibody (Molecular Probes, 1:500) and Alexa Fluor 488 goat anti-rat IgG antibody (Molecular Probes, 1:500) were used as secondary antibodies. Nuclei were counterstained with DAPI (Molecular Probes). These specimens were observed by a confocal microscope (A1R, Plan Apo λ 20×/0.75 NA, Nikon). The number of Parvalbumin7- and GCaMP6s-positive cells was counted by automatically detecting somata based on fluorescence signal intensity and size in confocal XYZ scan images using IMARIS (Zeiss, spot function). False-positive and false-negative cells were manually corrected.

### Calcium imaging of the cerebellum

For calcium imaging, a single-photon confocal microscope (A1R, Nikon, approximately 60 fps) or wide-field microscope (FN-1, Nikon, 16.7, or 33.3 fps) equipped with a CMOS camera (ORCA- Flash4.0) was used with a 40×/0.8 NA water-immersion lens. To extract changes in the fluorescence intensity, regions of interest (ROIs) were manually set using NIS-Elements (Nikon). Normalized changes in fluorescence intensity in the target regions (ROIs) were quantitatively analyzed by calculating ⊿F/F_0_ in the ROIs (⊿F/F_0_=[F(t)–F_0_]/F_0_, where F(t) and F_0_ are the fluorescence intensities at a certain time point and baseline time points, respectively). The baseline window for calculating F_0_ was adjusted according to the imaging frame rate and recording conditions (8–15 frames) but was kept constant within each analysis. Images were aligned as required (Fiji, TurboReg plugin). Resting conditions were defined as recording periods without externally presented visual stimuli. During these recordings, larvae were mounted in the same manner as in visual stimulation experiments, but no moving stripe patterns were presented.

### Characterization of responses and position of Purkinje cells

To characterize the Purkinje cell responses to visual stimuli, they were classified into four types based on the responses immediately after the start and end of the visual stimuli. Positive responses were defined as those with an onset within 2 s of the start or stop of the stripe movement and a response size (⊿F/F₀) of more than 12%, which was determined empirically.

For a detailed observation of GCaMP6s signals and identification of the recorded planes, XYZ confocal scanned images of the entire cerebellum were obtained after recording. Based on these images, the relative position of Purkinje cells in the cerebellum was calculated. Three-dimensional images were obtained using IMARIS (Zeiss).

### Detection of activated Purkinje cell populations

Purkinje cell populations that showed coordinated activation in wide-field calcium imaging were detected by the following processes using NIS-Elements AR (NIKON). First, activated Purkinje cell populations at each time point were extracted for each frame, with a value empirically set to ensure detected objects did not extend outside the cerebellum (⊿F/F_0_: 0.51––1.90%, size: 18- 388 pix, circularity: 0.2–0.7). To analyze the transitions of the activated Purkinje cell populations, a time-series tracking analysis of all detected objects across the recorded period was performed based on their position and speed. A series of objects that were considered to be identical across multiple frames was defined as “Purkinje cell clusters.” To visualize time-series changes in collectively activated Purkinje cell populations (objects), object contours were extracted from each frame, and contours from 100 frames were stacked using OpenCV. The stacked contours were colored red, blue, and green in chronological order, and further stacked images were created (Fig. 4g). To analyze the overlapped area among the collectively activated Purkinje cell populations, the maximum occupied area (the outer boundary of stacked contours) was generated every 10 frames. The overlapped area was calculated (Fig. 4h).

### Correlation analysis of Purkinje cell pairs

To assess the similarity in activity patterns between Purkinje cell pairs, Pearson’s correlations were calculated using *Δ* F/F_0_ data for approximately two minutes, excluding the periods of body movements. To obtain network diagrams, the analysis window was set to 30 s, and the correlation was measured every 15 s. To evaluate the coupling dynamics of the Purkinje cell pairs, cross-correlation analysis was performed on 120 s of calcium imaging data. The analyses were performed using custom-written Python or OriginPro (Lightstone).

### Laser ablation of inferior olive neurons

Laser ablation of the inferior olive via two-photon excitation was performed as described previously ^25^. In brief, the laser beam (λ = 780 nm, Δt ∼70 fs, Chameleon, SpectraPhysics) for two-photon excitation was focused on the inferior olive region of the agarose-embedded larva (*Tg(aldoca: GCaMP6s;hspzGFFgDMC28C; UAS: RFP)*, 6 dpf) on an inverted microscope (A1R MP+, Nikon) through an objective lens (Nikon Plan Apoλ 40x / 0.95 NA). The target area was determined by RFP signals of the IO neurons, and the stimulus ROIs were set in the IO regions. Laser irradiation was repeated until the tissue showed morphological defects and the RFP signals of the targeted IO neurons were absent (power: 60%, total irradiation time: 40–240 s/plane). By irradiating two to three different focal planes along the dorsoventral axis of the zebrafish, the entire region containing the inferior olive was ablated. After the ablation, the larvae were incubated at 28 °C before confocal microscopy imaging.

### Single-cell labeling of Purkinje cells

Plasmid DNA (15 or 7.5 ng/mL) *pT2K-aldoca-gap43-venus* (a generous gift from Prof. Masahiko Hibi, Nagoya University) was injected into 1-8-cell stage embryos. Venus signals in the cerebellum were observed under a confocal microscope (Olympus FV1000) at 6 dpf. Venus signals in single-cell-labeled Purkinje cells were extracted by binarization of confocal stack images, and their areas were measured (NIS-Elements).

### Enucleation of zebrafish larvae

Larvae anesthetized with 0.02% tricaine were mounted on 2% low-melting-point agarose. Bilateral enucleation was performed using a glass needle (2 dpf) or surgical scalpel (5 dpf). Post-surgery, larvae were reared in 1/3 Ringer’s solution until they were assayed at 6 dpf.

### Quantification and statistical analysis

Statistical analyses were performed using OriginPro (LightStone Co., Tokyo, Japan), the Statcel3 program (Bell Curve), and Python. Statistically significance was defined as P < 0.05. For two- group comparisons, the two-sample *t*-test, paired two-sample *t*-test, Mann–Whitney U test, or Kolmogorov–Smirnov test were applied. The Tukey–Kramer test was used for multiple comparisons.

## Notes

### Competing Interest Statement

The authors have declared no competing interest.

### Summary of Updates

Edit the author list, text, and Figures.

